# The DMT and Psilocin Treatment Changes CD11b+ Activated Microglia Immunological Phenotype

**DOI:** 10.1101/2021.03.07.434103

**Authors:** Urszula Kozłowska, Aleksandra Klimczak, Kalina Wiatr, Maciej Figiel

## Abstract

Psychedelics are new, promising candidate molecules for clinical use in psychiatric disorders such as Treatment-Resistant Depression (TRD) and Post Traumatic Stress Disorder (PTSD). They were recently also proposed as molecules supporting neural tissue repair by anti-inflammatory properties. Here we reported that two classic psychedelics, DMT and psilocin, can influence microglial functions by reducing the level of TLR4, p65, CD80 proteins, which are markers of the immune response, and upregulat TREM2 neuroprotective receptor. Psilocin also secured neuronal survival in the neuron-microglia co-culture model by attenuating the phagocytic function of microglia. We conclude that DMT and psilocin regulate the immunomodulatory potential of microglia. Of note, psychedelics were previously reported as a relatively safe treatment approach. The demonstrated regulation of inflammatory molecules and microglia phagocytosis suggests that psychedelics or their analogs are candidates in the therapy of neurological disorders where microglia and inflammation significantly contribute to pathogenic disease mechanisms.

## Introduction

Psychedelic therapy is undoubtedly a recent breakthrough in psychiatry for treatment-resistant depression, post-traumatic stress disorder (Nutt et al., 2020). Recent studies in animal models suggest that psychedelics may also display a broader spectrum of applications in neuronal damage and neuropsychiatric conditions such as Traumatic Brain Injury (TBI), Alzheimer’s Disease (AD), and others (Scott and Carhart-Harris, 2019; Vann Jones and O’Kelly, 2020). Their therapeutic effects are related to immunomodulatory potential, ability to induce neurogenesis, and neural plasticity; however exact mechanisms are yet to be identified (Flanagan and Nichols, 2018; Ly et al., 2018).

The reported anti-inflammatory potential of psychedelics is appealing since the state of acute brain inflammation is the direct cause of neural cell death and limitation of the regeneration process. Such conditions are frequently observed in neurodegenerative diseases or Traumatic Brain Injury (Geloso et al., 2017), (Loane and Kumar, 2016). Anti-inflammatory drugs are recently proposed as a therapeutic strategy since they can reduce microglia’s harmful activity (Jarrott and Williams, 2016). However, anti-inflammatory pharmacologic approaches used in the clinical treatment so far may cause serious adverse effects related to broad immune system suppression.

Microglia is a particular type of brain-specific macrophages, which display a broad spectrum of actions to support neuronal network rearrangement, neural tissue regeneration, and protection from pathogens (Ikegami et al., 2019; Diaz-Aparicio et al., 2020). In the healthy brain, the microglial cells work in favor of tissue rearrangement and regeneration, induction of NPC neurogenesis, and control of synaptic pruning directly or indirectly by recruiting astrocytes. The phagocytosis activity of microglia serves for cleansing the local environment from pathogens, damaged or infected cells, cellular debris, and toxic protein aggregates (Diaz-Aparicio et al., 2020; Matejuk and Ransohoff, 2020). Like macrophages, the microglia infiltrates the local environment detecting the infection or tissue damage signals through Pattern Recognition Receptors (PRRs), and can present the antigen to T-cells. During infection, microglia responds to bacterial LPS via Toll-like Receptor 4 (TLR4), starting the innate immune response mediated by NF-κβ (p65) activation. The NF-κβ regulates pro-inflammatory protein secretion by microglia, which may attract other immune cells to cross the Blood-Brain Barrier (Kigerl et al., 2014) (Dionisio-Santos et al., 2019). The activated microglia upregulates CD40, CD80, or CD86 co-stimulatory molecules expression, which provides the second signal for antigen presentation via MHC to T-cells (Sharpe, 2009; Nayak et al., 2014). If the process of microglia immune-regulation excessively amplifies, it may lead to the attraction of unwanted immune cells, the expansion of microglial phagocytosis to functional neurons, and other pro-inflammatory actions which leads to acute tissue damage (Janda et al., 2018; Yanuck, 2019) (Mohebiany et al., 2020).

Another important regulator of several microglia functions is the Triggering Receptor Expressed on Myeloid Cells 2 (TREM2), which plays an essential role in proper microglial phagocytosis (Deczkowska et al., 2020). TREM2 reacts to anionic molecules, like Danger associated molecular patterns (DAMPs), such as toxic protein aggregates and Pathogen associated molecular patterns (PAMPs), including bacterial lipopolysaccharide (LPS) (Gyengesi and Münch, 2020). Activation of TREM2 initiates anti-inflammatory pathways in microglia; therefore, dysregulation of its expression may lead to neurodegenerative diseases, where it may be a promising therapeutic target (Deczkowska et al., 2020).

In the current study, we aimed to investigate whether the application of classic psychedelics, such as DMT and psilocin, could be beneficial for neural tissue homeostasis and the promotion of pro-regenerative and anti-inflammatory features. We tested DMT and psilocin’s influence on mouse primary CD11b+ microglia by measuring the protein expression of factors crucial for innate and adaptive immune response regulation (TLR-4, NF-κβ, CD80). We also measured the changes in TREM2 protein expression, which is recently studied in the context of neuroprotection and proper microglial phagocytosis. Finally, we checked if DMT and psilocin affect microglial phagocytosis of healthy neurons after LPS stimulation. Our investigation examines classic psychedelics’ influence on the model of activated microglia also together in neurons. Therefore, we mimic the therapeutic influence of psychedelics in neurological disorders where inflammation and microglia significantly contribute to pathogenic disease mechanisms.

## Materials and Methods

### Microglia culture

DBA1 mice age between 3 to 6 months were euthanized according to local ethical law regulations. Shortly after death (up to 1 min), the bodies underwent full-body perfusion with ice-cold PBS. After the brain extraction, mixed glial cell culture from each mouse was isolated by homogenizing the brain in a glass homogenizer (Sigma) and then purified by mixing 17 mL PBS brain lysate with 4,5 mL Percoll Plus (Sigma, Saint Louis, Missouri, USA). After 15 min 500g centrifugation in +4 □C, the pellet was resuspended in DMEM/F12 (Thermo Fisher, Waltham, Massachusetts, USA), 10% Fetal Bovine Serum (Gibco, Origin: Brazil, Campinas, Brazil), 1% penicillin/streptomycin, 1% L-glutamine (Sigma-Aldrich, Steinheim, Germany), 1.5 µg/mL, sheep wool cholesterol, 2 ng/mL TGF-β, 15 ng/mL IL-34 and 20 ng/mL bFGF (Sigma, Saint Louis, Missouri, USA). The culture media was changed every 4 days until cell cultures reached full confluence. The ratio of astrocytes to microglia was then assessed with immunofluorescence (IF) and flow cytometry (FCM) techniques. In order to activate the microglia with LPS, after reaching the monolayer, the culture at P0 was washed 2 times with PBS and incubated 24h in growth factor-free DMEM/F12 media, supplemented with 1% penicillin/streptomycin, 1% L-glutamine, and 1.5 µg/mL sheep wool cholesterol (VEH). After 24h incubation in VEH media, new VEH media containing 150 ng/mL LPS (Sigma, Saint Louis, Missouri, USA) was added for 24h (Fig. 2). For detaching, the cells were washed twice with PBS and then digested with pre-warmed TrypLE (Thermo Fisher, Waltham, Massachusetts, USA) for 10 min in +37 □C. After incubation, the media was neutralized 1:1 with culture media, and the cells that remained attached were taken off by gentle but dynamic pipetting. The cells were then centrifuged 470 g for 5 min in +4 □C.

**Fig 1:**
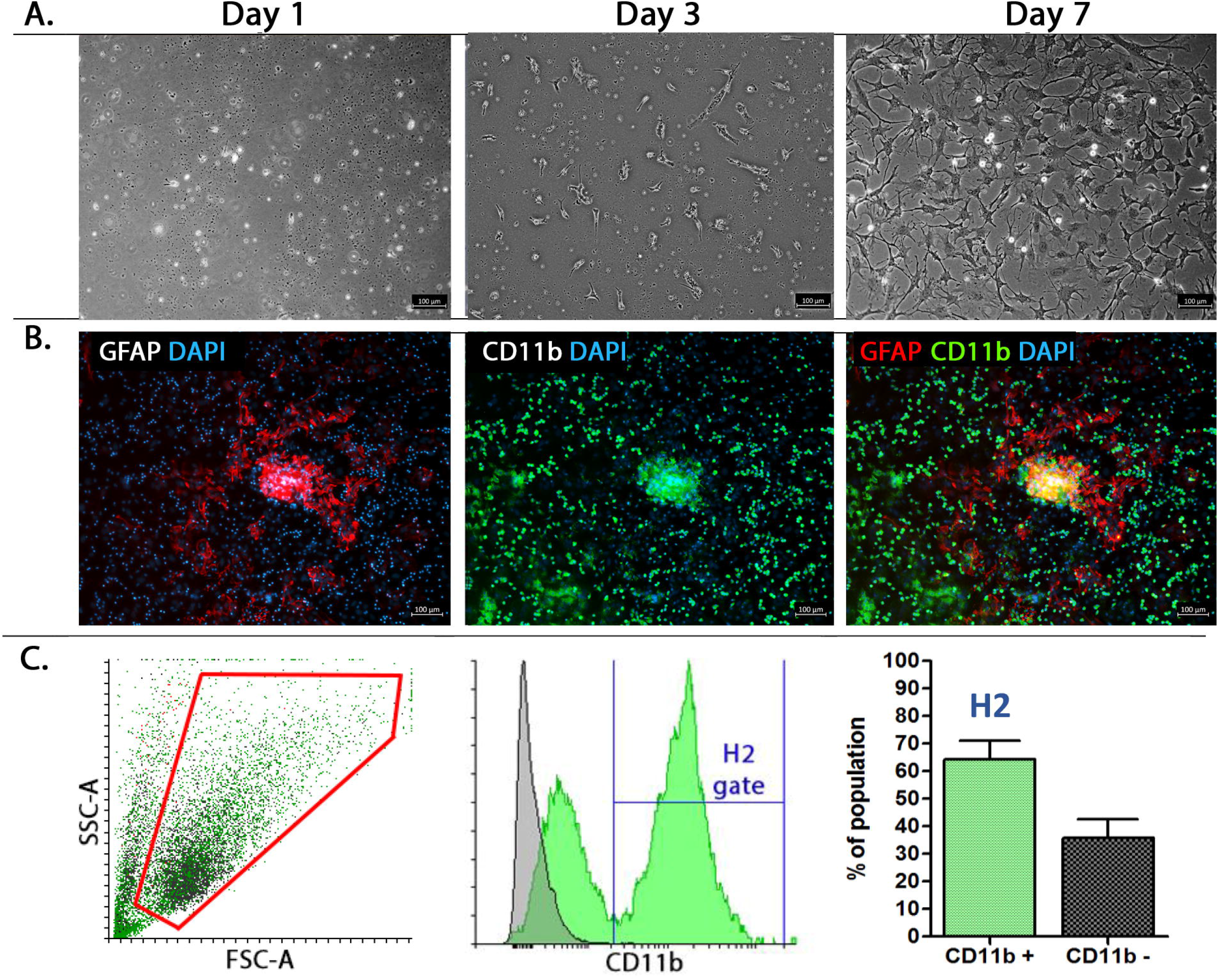
Characteristics of the cells isolated from 3 – 6 months old DBA/1 mice brain: A) After 7 days, the cells formed numerous, dense colonies. B) The GFAP and CD11b staining revealed a mixed population of astrocytes and microglia cells. C) FCM analysis revealed that 64.3% of the population in mean express CD11b marker (SEM = 1.96), error bars: SEM; total number of samples n□=□12 per experimental group (grey – unstained control, green – CD11b. Scale bar – 100 µm.

**Fig 2:**
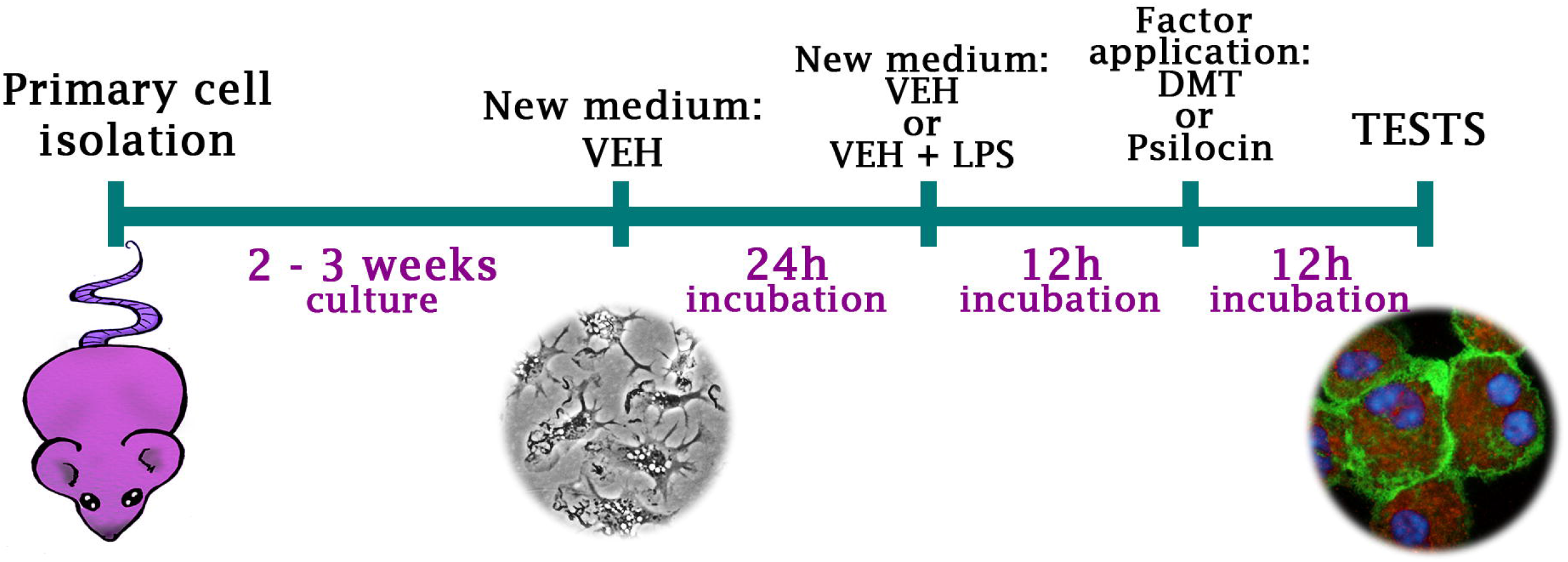
Design of the microglia phenotype experiment: The primary heterogeneous population was isolated from DBA mice brain (n=30, number of T-25 flasks =60), and cultured 2-3 weeks until monolayer. Then the microglia culture media was changed for VEH media for 24h. Next, the VEH media was changed for VEH or VEH + 150 ng/mL LPS media. After 12h of incubation, the DMT or Psilocin was added to reach a final concentration of 100 µM (research groups) or untreated (control group) for another 12h. The tests were performed immediately after all the incubations.

### Neurospheres culture

The neurospheres were isolated from C57BL/6 mouse E10 embryos according to the modified protocol described in Ebert 2013 (Ebert et al., 2013). In brief, embryonic brains were dissociated by trypsin (Merck, Darmstadt, Germany), washed 2 times in HBSS (Thermo Fisher Scientific, Waltham, MA, USA), supplemented with 1x penicillin-streptomycin (Thermo Fisher Scientific, Waltham, MA, USA), and pre-plated on uncoated plates for 2 hours. The neurospheres were then cultured on non-adherent 6-well plates in neurosphere medium containing: 7:3 mixture of DMEM with Hams F12 Nutrient mix supplemented with 2% of B27 supplement, 100 ng/mL EGF, 100 ng/mL bFGF (Thermo Fisher, Waltham, Massachusetts, USA), 5 μg/mL heparin (Sigma, Saint Louis, Missouri, USA) and 1% penicillin/streptomycin, 1% L-glutamine (Sigma, Saint Louis, Missouri, USA). The maintenance medium was changed every 2-3 days, and passages were performed every 4–5 days using mechanical disassociation (Svendsen et al., 1998).

### NSC differentiation

Neurospheres were dissociated to single-cell and were seeded on poly-L-ornithine/laminin (Sigma, Saint Louis, Missouri, USA) coated plates for 24h in neurosphere medium. After the cells fully attached to the surface, the medium was changed to neuronal differentiation medium: Neurobasal and DMEM/F12 1:1, 1% of B27 supplement, 30 ng/mL GDNF, 50 ng/mL BDNF (Thermo Fisher, Waltham, Massachusetts, USA), 1% penicillin/streptomycin, 1% L-glutamine (Fig. S1).

### Immunofluorescent staining

The cells attached to the 48-well plate were fixed for 10 min with 10% buffered formalin Plus (Sigma, Saint Louis, Missouri, USA) and then permeabilized for 15 min with 0.01% Tween-20 (Serva, Heidelberg, Germany). The cells were then incubated in 1% BSA (Abcam Cambridge, Great Britain) and 10% Goat Serum (Thermo Fisher, Waltham, Massachusetts, USA) for 1h RT. Then the primary antibody solution was added and incubated (GFAP, Thermo Fisher, 1:200 4h, RT), (TREM2, Bioss, Woburn, MA, USA 1:200, overnight, +4□C), (Nf-κβ, Invitrogen, Carlsbad, CA, USA 1:200, overnight, +4 □C), (TLR4, Bioss, 1:200, overnight, +4□C), (CD11b, Bio-Rad, Hercules, CA, USA, 1:100, overnight, +4□C). After incubation, the primary antibody was rinsed off with PBS, and the secondary antibody (Alexa Fluor 488nm or 594nm, Thermo Fisher, Waltham, Massachusetts, USA) was applied in a concentration of 1:400 (488nm) or 1:200 (594nm) for 1h in RT. After rinsing off three times with PBS the secondary antibody solution, the cell nuclei were visualized with DAPI Sigma, Saint Louis, Missouri, USA staining.

### Immunofluorescence analysis

The stained cells were analyzed with Zeiss Axio Observer 2 microscope, using Alexa Fluor 488 nm settings for CD11b staining: 60% blue laser strength, 1000 ms acquisition, Alexa Fluor 594 nm settings for TLR4, TREM2, and Nf-κβ stainings: 100% orange laser strength, 5000 ms acquisition. The optimal acquisition time and laser strength were assessed as compared to negative control staining. The image analysis was adjusted in ZEN 2.0 software, setting 1050 units lower - 3500 units upper curve threshold for TLR4 and TREM2 staining, and 1050 units lower - 4000 units upper curve threshold for Nf-κβ staining. The Corrected Total Cell Fluorescence (CTCF) was measured using ImageJ Software and calculated using equation **CTCF = D**_**n**_ **– (A**_**n**_ **x B**_**n**_), where **D**-integrated density, **A**-area of the selected cell, **B**-mean background, **n**-event number (Fig. S2).

### Flow cytometry

The cells (dethatched by TrypLE digestion as previously described) were incubated for 10 min in 1% BSA (Abcam Cambridge, Great Britain) containing PBS to block unspecific reactions. Then the cells were centrifuged 470 g, 5 min, +4 □C, and resuspended in PBS. The cells were divided into two fractions – one fraction dedicated for compatible isotype control stainings (Thermo Fisher, Waltham, Massachusetts, USA) and the second fraction for CD11b (Bio-Rad, Hercules, CA, USA) staining (1 h on ice) and secondary antibody (Alexa Fluor 488 nm, 1:400 for 30 min on ice). After rinsing in ice-cold PBS and +4 □C 470 g centrifugation, the cells were stained for 30 min with 2 µL of Thermo Fisher antibodies (Armenian Hamster IgG, CD80, PE), (Rat IgG2, CD11c, APC), (Armenian Hamster IgG CD163, APC). Each test was performed with density 1× 10^5^ cells in 150 µL. The reaction was ended by adding 2 mL of PBS into the probes, and the cells were 10 min centrifuged in +4 □C with 470 g speed. The measurements were made with BD Fortessa Cytometer in DIVA software after proper compensation. Before measurements, DAPI was added into cell suspension as a control to gate out the dead cells.

### PKH cell staining

The cells collected from the culture plate were stained with PKH (Sigma, Saint Louis, Missouri, USA) by 5 min incubation in a mixture containing 2 µL PKH and 500 µL diluent C on the ice. Subsequently, the dye was neutralized by adding 10 mL of PBS containing 1% BSA and centrifuged 470 g in +4 □C for 5 min. The cells were again washed with 5 mL 1% BSA in PBS.

### Neural-glial co-cultures

The neural cells were differentiated in 24-well plates for 5 days. The microglia cells (LPS-stimulated and non-stimulated) from monolayers were detached from culture flasks, PKH67 stained, and added to neural cultures (5 × 10^4^ per well). The heterogeneous populations of cells isolated from DBA mice brains were 24h pre-treated with LPS or non-treated (VEH). After 24h, cells were taken off the culture flask, stained with PKH67, and added into wells with neural cells to form co-cultures, and observed for 24h in six experimental variants: 1) 100 µM of DMT in VEH media, 2) 100 µM of DMT + 150 ng/mL LPS, 3) 100 µM of psilocin in VEH media, 4) 100 µM of psilocin + 150 ng/mL LPS, 5) control (VEH), 6) control with 150 ng/mL of LPS. In the LPS treated groups, cells were pre-treated with 150 ng/mL LPS for 24h before initiation of the experiment. The continuous observation was done for 24h using a Zeiss Axio Observer microscope in 10 min intervals.

### Analysis of cell morphology

The microglia cells morphology was measured from contrast-phase pictures using ImageJ software. The branch length was measured using the skeletonization method, and fractal dimension, lacunarity, span ratio, and circularity were measured using FracLac plug-in, both described in the protocol by (Young and Morrison, 2018).

### Statistical analysis

The experiments were made at least three times in duplicates. Statistical analysis was done using Graph Pad Prism 5. All *p*-values were calculated by using a t-test for unpaired samples with Welch’s correction. The graphs are presented as mean values with a standard error of the mean (SEM).

## Results

### DMT and psilocin modify the morphology of microglia

The primary microglial cells were isolated from DBA mice brains as a heterogeneous population. The heterogenous population was evaluated by estimating the percent of CD11b+ microglia vs CD11b negative cells. After reaching the monolayer in 3 weeks, an average of 65.3% of cells expressed marker CD11b+ as analyzed by FCM (Fig. 1). The staining for GFAP also revealed the presence of numerous astrocytes in the population (Fig. 1).

After reaching the monolayer, to limit external stimuli, microglia cultures were pre-treated for 24h in serum-free defined media in which all grow factors (except cholesterol) were omitted (VEH media). The microglia cultures were either preincubated for 24h in media supplemented with 150 ng/mL LPS or cultured in fresh VEH media. The cultures with LPS and without LPS were subsequently treated for 12h with 100 µM DMT or 100 µM psilocin or left untreated (Fig. 2). There were subtle changes in microglia morphology in the VEH group (Fig. 3A). Cells cultured in VEH media stayed mostly in the rod/bipolar or small, round shape; however, in the group treated with psilocin and DMT, long, ramified cells with long branches were often observed (Fig 3B, D). After 24h incubation with LPS, the microglia were rather round and acquire a ramnified morphology. However, after 12h of DMT or psilocin treatment, many cells acquired ameboid morphology (Fig. 3C). DMT or psilocin-treated cells in comparison to control (just LPS-treated group) were characterized with lower span ratio (**p<0.0063 DMT),(**p<0.0098 psilocin), higher circularity (*** p<0.0003 DMT), (**p<0.001-psilocin) and lower fractal dimension (** p < 0.0083 DMT) (Fig. 3 E, F, G).

**Fig 3:**
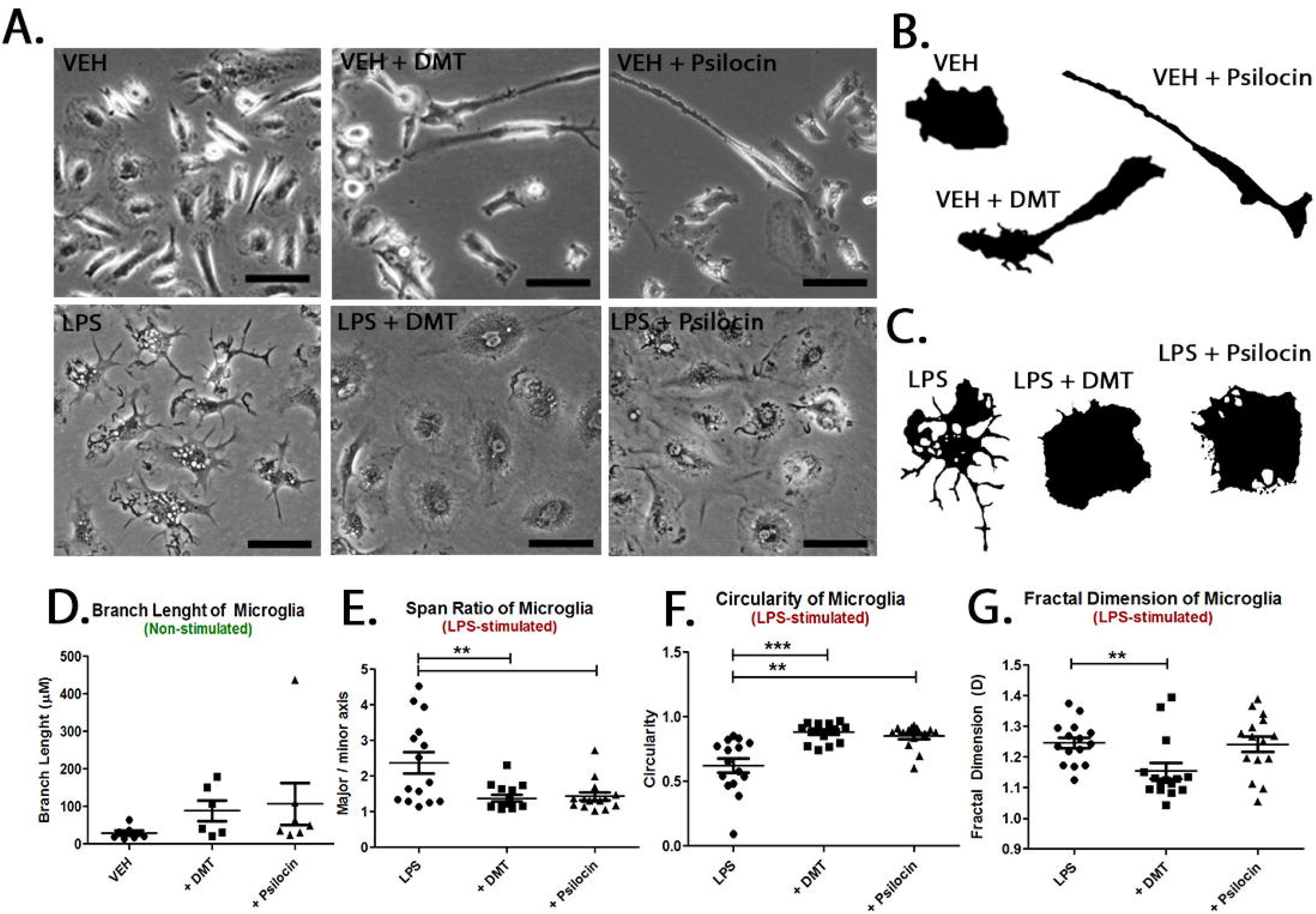
The LPS, DMT, and psilocin treatment altered the morphology of CD11b+ microglia cells: A) The contrast-phase pictures of microglia morphology in VEH (upper panel) and LPS (lower panel) groups. Some of the cells in the VEH group were characterized with longer branches after 12h of DMT and psilocin treatment. In the LPS group, DMT and psilocin treatment resulted in reassuming ameboid morphology as compared to just LPS-treated cells, which morphology was ramified. B) Different forms of microglia morphology in VEH group, C) Different forms of microglia morphology in LPS group, D) branch length of microglia in VEH group (VEH – DMT p, <0.0893), (VEH-psilocin, p<0.2148), E) span ratio of microglia in LPS group (LPS – DMT **p<0.0063), (LPS-psilocin, **p<0.0098), F) circularity of microglia in LPS group (LPS-DMT, *** p<0.0003), (LPS-psilocin, **p<0.001), G) fractal dimension of microglia in LPS group (LPS-DMT** p < 0.0083), (LPS-psilocin, p <0.8911 Unpaired Student’s t-test; error bars: SEM; the total number of biological replicates in VEH group: n = 3, the total number of events: n = 556, events per variant: n= 82 – 284. Total number of biological replicates in LPS group: n = 3, total number of events: n = 45, events per variant: n= 15, Scale bar= 50 µm.

### The TLR4 protein level is downregulated in the psychedelic-treated group after LPS-stimulation

To investigate how DMT and psilocin affect microglia immunological phenotype, we studied the expression of markers involved in inflammatory pathways: TLR4, Nf-κβ, and CD80; and TREM2, which is suggested to be involved in neuroprotection and regulation of microglial phagocytosis. We measured TLR4, TREM2, and Nf-κβ (p65) expression using the CTCF analysis of CD11b+ microglia.

TLR4 receptor binds LPS which activates pro-inflammatory cascade, leading to upregulation of co-stimulatory molecules expression in antigen-presenting cells (APC) such as microglia. We observed that the expression of TLR4 did not significantly change in the VEH group after the application of both psychedelics. However, after LPS stimulation, the TLR4 expression on CD11b+ cells measured by fluorescence intensity was significantly lower in groups stimulated with DMT and psilocin (***p < 0.0001) as compared to the control group (Fig. 4, Fig. S3).

**Fig 4:**
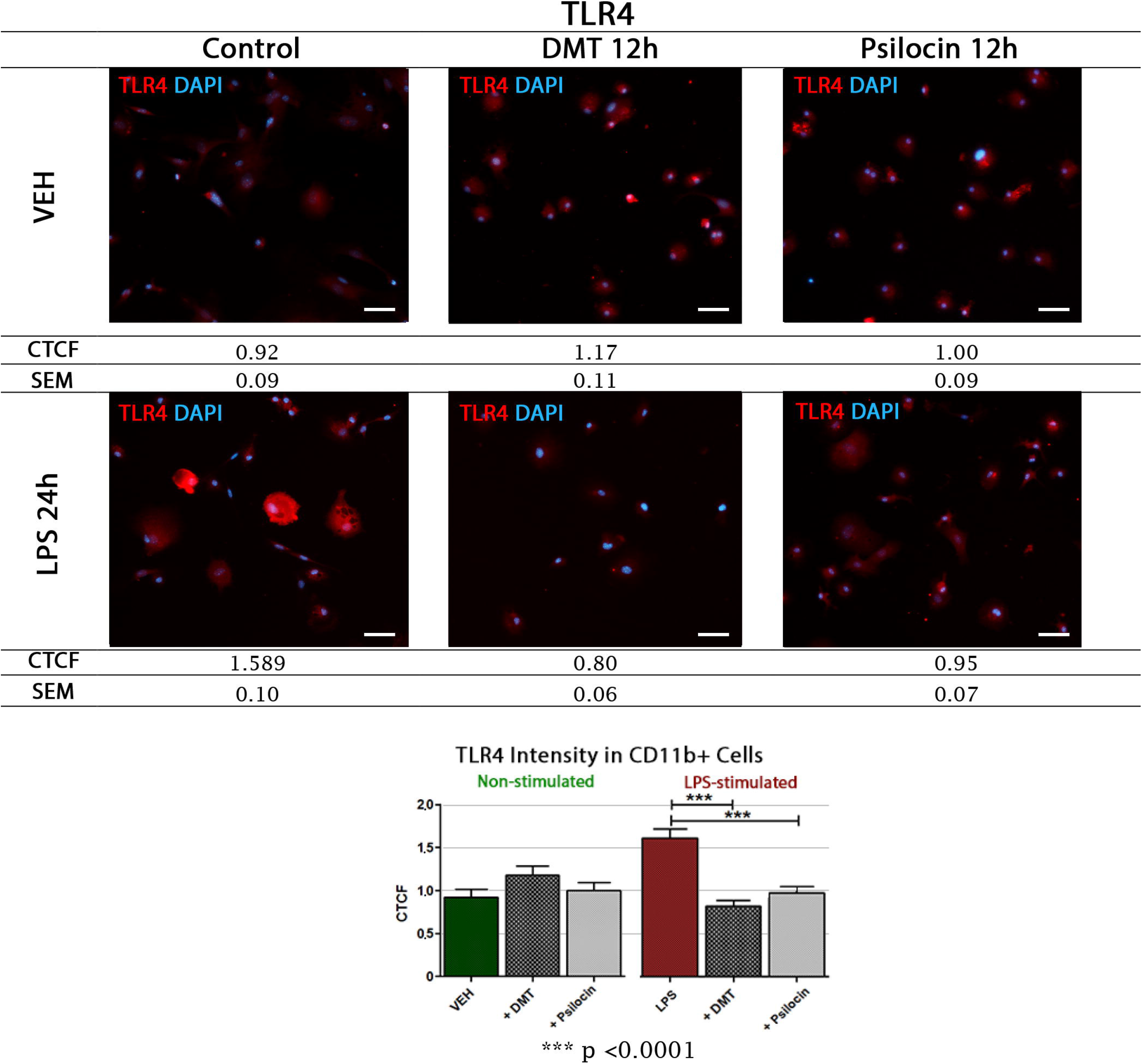
The DMT and psilocin 12h stimulation decreased fluorescence intensity of TLR4 on CD11b+ microglia cells after LPS treatment: The immunofluorescent staining (IF) revealed that in the VEH group, the incubation with DMT and psilocin did not change the TLR4 expression. However, after LPS treatment, the application of DMT and psilocin significantly (***p < 0.0001) decreased TLR4 fluorescence intensity, which could be comparable with the intensity measured in the VEH group. For double staining pictures, please see supplementary data.. Unpaired Student’s t-test; error bars: SEM; the total number of biological replicates in VEH group: n = 6, the total number of events: n = 418, events per variant: n= 131 – 153. Total number of biological replicates in LPS group: n = 6, total number of events: n = 437, events per variant: n= 143 – 150 Scale bar – 50 µm.

### DMT and psilocin downregulate Nf-κβ p65 protein level in both control and LPS conditions

The signaling cascade from LPS – stimulated TLR4 receptor under normal conditions results in activation of Nf-κβ (p65), a transcription factor that activates the expression of the pro-inflammatory genes (Liu et al., 2017). The microglial Nf-κβ (p65) fluorescence intensity decreased after DMT and psilocin stimulation both in VEH, and LPS-treated groups (** p < 0.002, ***p < 0.0001). In the picture (Fig. 5), which presents DMT-treated cells in the VEH group, the cells tagged with yellow arrows are the CD11b+ microglia cells (for the double-stained picture, see Fig. S4).

**Fig 5:**
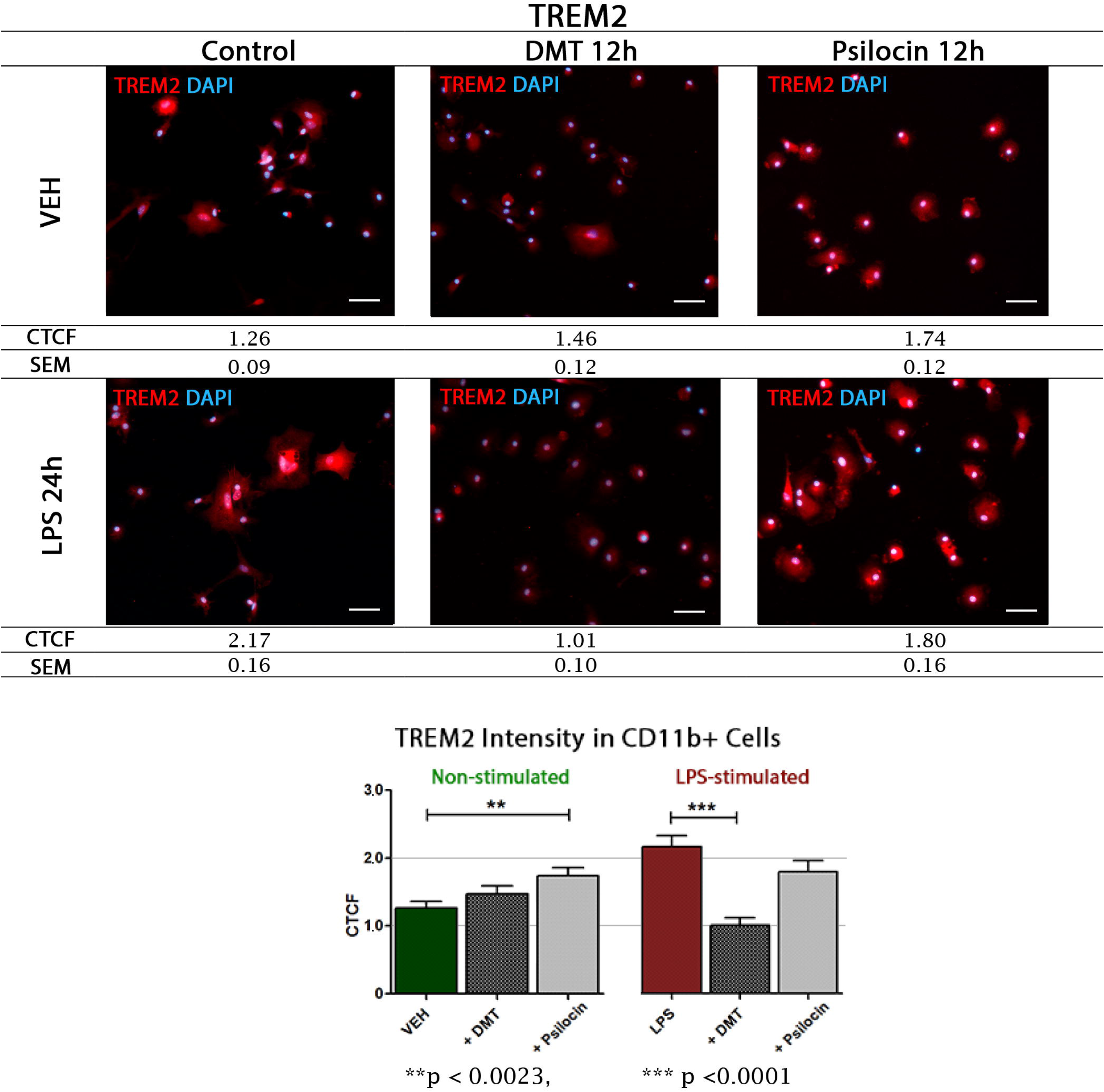
DMT and psilocin downregulated the expression of NF-κβ in CD11b+ microglia cells unstimulated and after LPS stimulation: The IF staining revealed that CD11b microglia loses the expression of NF-κβ in VEH and in LPS-stimulated groups when treated both with DMT (** p < 0.002 VEH group, *** p < 0.000 LPS-stimulated group) and psilocin (*** p < 0.001 VEH and LPS-stimulated groups). For double staining pictures, please see supplementary data. Unpaired Student’s t-test; error bars: SEM; total number of samples in VEH group: n = 6, total number of events: n = 410, events per variant: n= 133 – 143. Total number of biological replicates in LPS group: n = 6, total number of events: n = 421, events per variant: n= 131 – 159. Scale bar – 50 µm

### DMT and psilocin downregulate CD80 co-stimulatory molecule

CD80 co-stimulatory molecule is necessary for antigen presentation via MHC T-cells, which is crucial for adaptive immune response development. Blocking the co-stimulatory molecule protein domains with specific factors for their inactivation is one of the immunosuppressive strategies called co-stimulatory blockade(Melvin et al., 2012; Maltzman and Turka, 2013). The FCM analysis of CD80 marker in CD11b+ (H2 gated) cells revealed that DMT and psilocin application to the culture media decreases CD80 co-stimulatory molecule expression on both VEH and LPS-treated microglia. In the untreated group, both DMT and psilocin decreased CD80 protein level; however, the statistical significance (*p > 0.0172) was observed only in the group where the psilocin was applied. In the LPS-treated microglia, the decrease of CD80 protein level in DMT and psilocin stimulated groups was greater than in the group where no psychedelics were applied. Statistical significance was calculated for both DMT (*p > 0.0115) and psilocin (**p > 0.0069) (Fig. 6).

**Fig 6:**
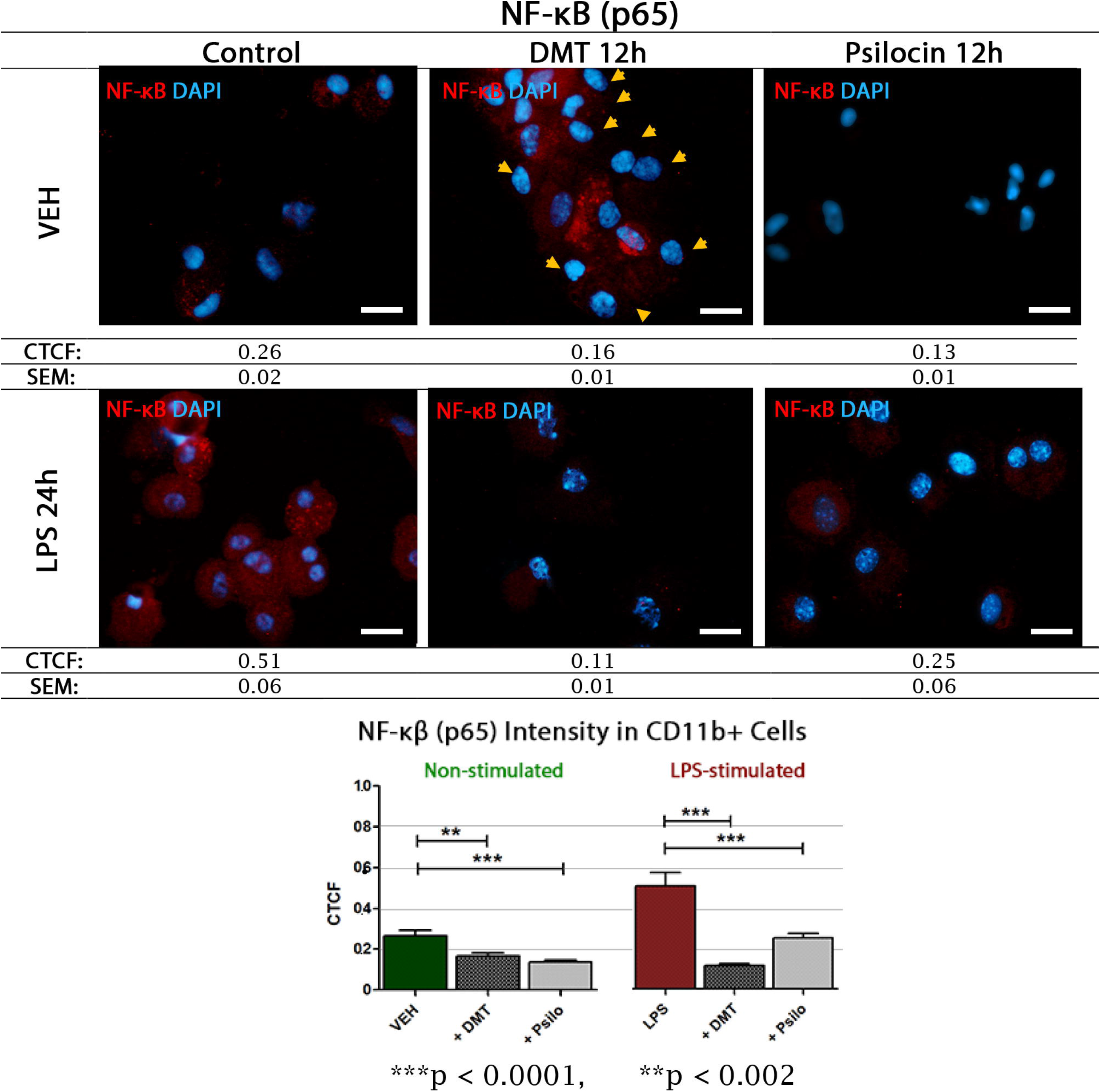
The observation of CD80 surface protein expression change in CD11b+ microglia after 100 µM DMT or 100 µM psilocin administration: In the VEH group the expression of CD80 decreased after 12h incubation with DMT (p = 0.0519) and psilocin (*p > 0.017). In the LPS-treated group the administration of agents for 12h, resulted in significant decrease of CD80 expression in CD11b+ microglia after DMT (*p > 0.0115) and psilocin (**p > 0.0069) treatment. Unpaired Student’s t-test; error bars: SEM; the total number of samples n□=□19, n□=□6-7 per experimental group).

### Psilocin but not DMT upregulates TREM2 protein level in the VEH group

TREM2 is a receptor present on myeloid cells, recently studied in the context of proper regulation of microglial phagocytosis and its neuroprotective properties (Deczkowska et al., 2020). The stimulation with psilocin, but not DMT significantly (**p < 0.0023), increased the expression of TREM2 in the VEH group. Interestingly, in the LPS-treated group, treatment with DMT significantly (*** p< 0.0001) decreased TREM2 fluorescence intensity. In contrast, psilocin administration did not cause this effect, leaving TREM2 expression on a level similar to the VEH control group (Fig. 7, Fig. S5).

**Fig 7:**
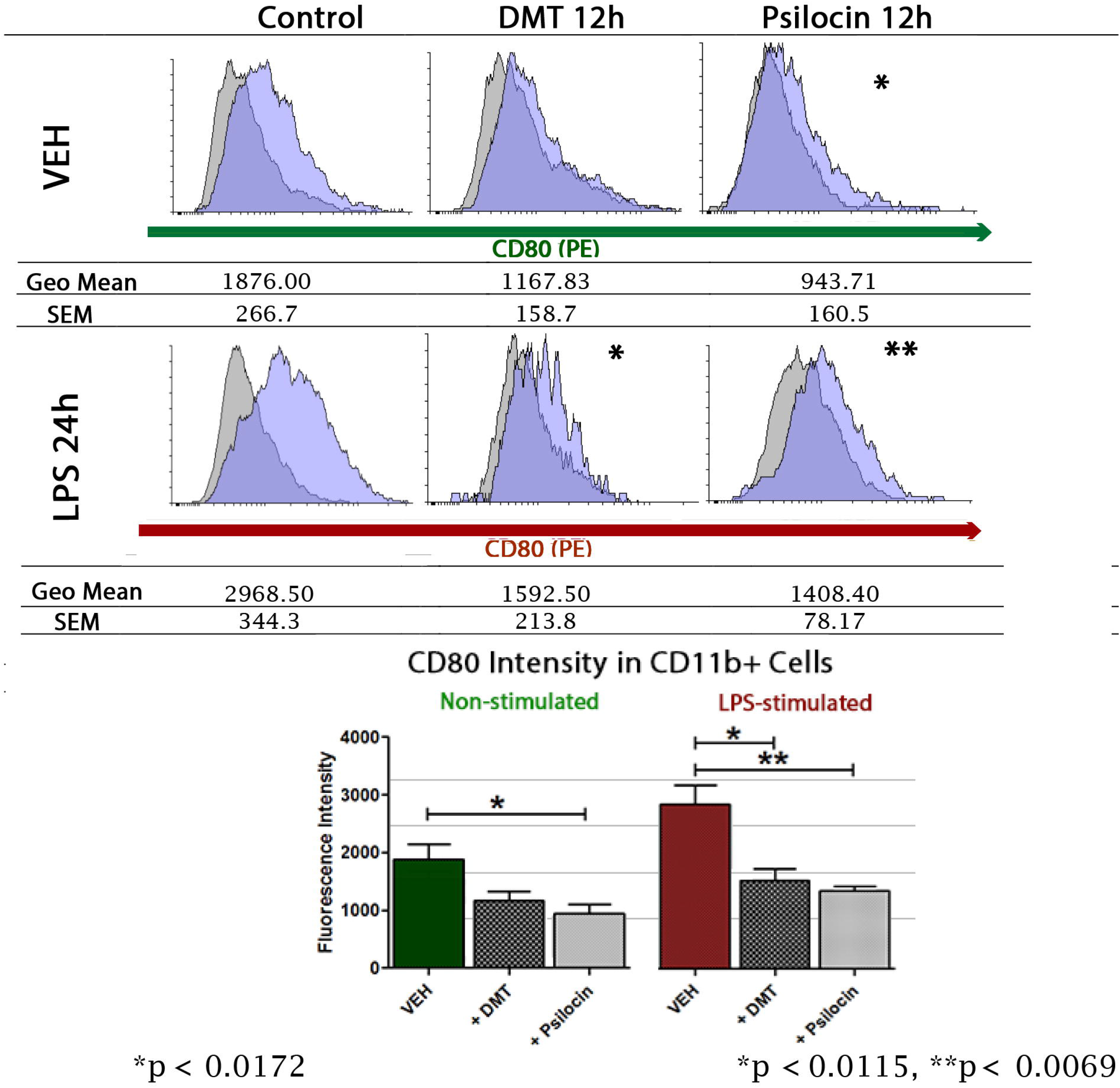
The DMT and psilocin 12h stimulation changes TREM2 fluorescence intensity on CD11b+ microglia cells: The IF staining revealed that the incubation with psilocin but not DMT increased the TREM2 expression (**p < 0.0023) in the VEH group. After LPS treatment, the application of DMT significantly decreased TREM2 expression (***p < 0.0001), whereas its expression was maintained in the psilocin group. For double staining pictures, please see supplementary data. Unpaired Student’s t-test; error bars: SEM; total number of biological replicates in VEH group: n = 6, total number of events: n = 444, events per variant: n= 138 – 158. Total number of biological replicates in LPS group: n = 6, total number of events: n = 435, events per variant: n= 142 – 151. Scale bar – 50 µm

### Psilocin but not DMT attenuates healthy neurons microglial phagocytosis

The microglia is essential in preserving tissue homeostasis, but sometimes pro-inflammatory and pro-regenerative microglia functions become imbalanced. Hyperactive microglia may react to compounds on the surface of stressed but healthy neurons and proceed with phagocytosis, seriously deteriorating the process of neural healing or even triggering neural damage (Brown and Neher, 2014). The study in co-cultures revealed that the application of 100 µM psilocin into culture media resulted in reduced phagocytosis of healthy neurons by microglia compared to DMT, both in VEH and LPS groups. This observation suggests that psilocin may display more robust neuroprotective properties than DMT by reducing healthy neuron phagocytosis by microglia (Fig. 8, see supplementary videos V1, V2, V3. V4).

**Fig 8:**
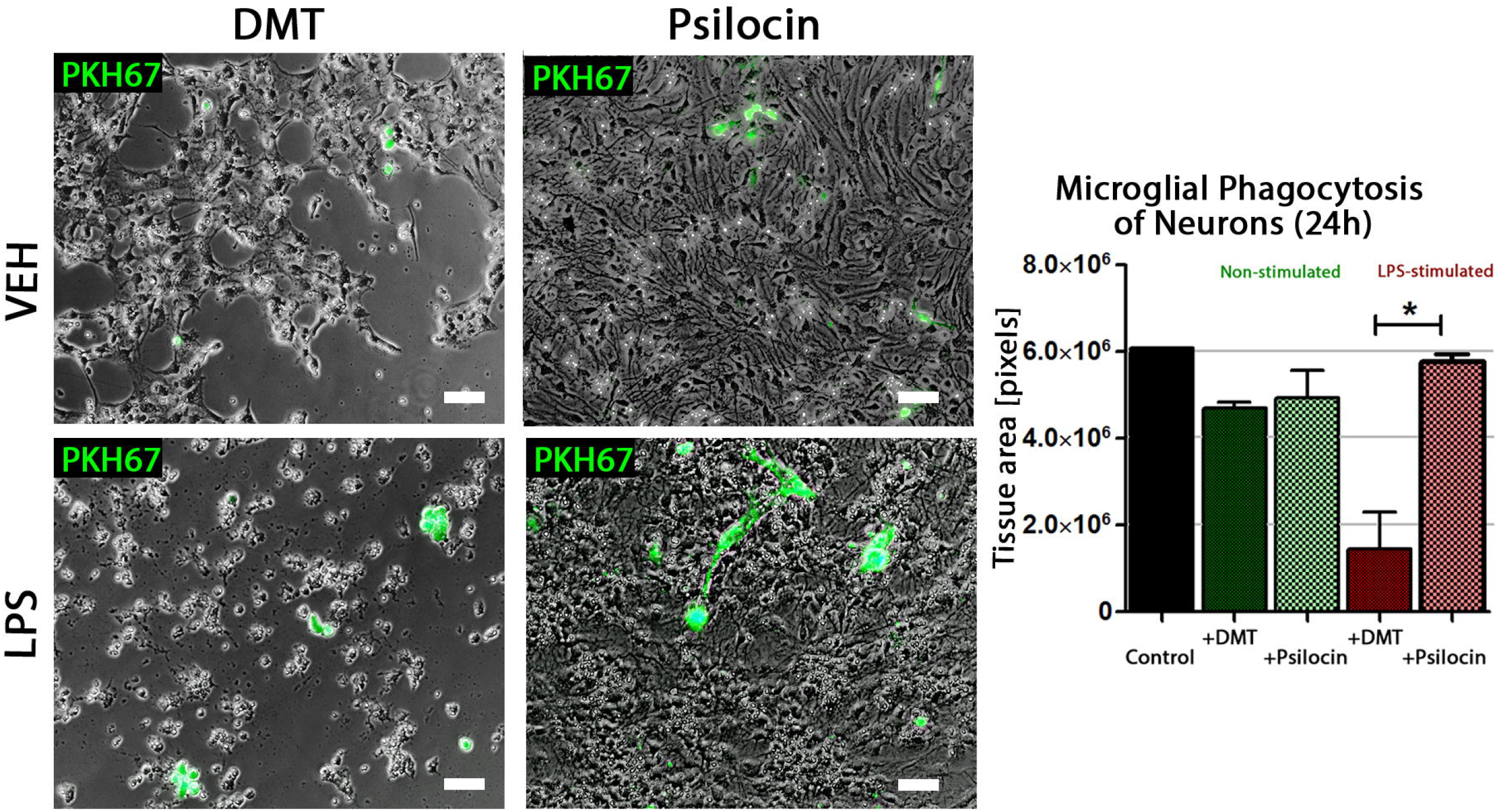
Co-cultures of neural cells and a heterogeneous fraction of adherent cells isolated from DBA mice brains with and without LPS stimulation: The phagocytosis of healthy neutral cells was observed in both VEH and LPS-treated group after 24h of co-culture, as compared to control of neural monoculture. 24h of co-culture after LPS-stimulation resulted in greater neural phagocytosis in DMT-group as compared with the psilocin group, where the damage was visibly smaller (*p<0.0158). Unpaired Student’s t-test; error bars: SEM; the total number of samples n□=□15, n□=□3-4 per experimental group. Scale bar – 50 µm

## Discussion

The inflammatory environment which limits brain regeneration can be caused by hyperactive microglial cells as often seen in neurodegenerative disorders and brain damage, such as ALS (Geloso et al., 2017), Alzheimer Disease (Hemonnot et al., 2019), and Parkinson Disease (Lull and Block, 2010), Multiple Sclerosis (Plastini et al., 2020) and TBI (Loane and Kumar, 2016). Moreover, stem cell grafts are often proposed as cell therapy for neurodegeneration and brain damage; however, host vs. graft immune response can be a severe limitation (Piquet et al., 2012). Current anti-inflammatory strategies that attenuate hyperactive immune response and microglia are recently proposed as a therapy for neurologic disorders. Unfortunately, therapeutics with anti-inflammatory properties such as antibiotic minocycline failed in the experimental treatment of AD patients and were not well tolerated at a higher dose (400 mg) (Howard et al., 2020). The limitation of minocycline and other anti-inflammatory drugs, whether steroid or non-steroid, is their broad-band influence on immune response and their severe adverse effects after long-term use (Davis and Robson, 2016). Many other novel therapeutics based on humanized antibodies targeting specific anti-inflammatory molecules are not suitable to crosse blood-brain barrier to treat brain inflammation (Klotz and Wiendl, 2013; Pardridge, 2020) The anti-inflammatory therapeutic strategy should be more selective and involve cytokine-suppressive anti-inflammatory drugs (CSAIDs), that would target pro-inflammatory cytokine production pathways in astrocytes and microglia (Gyengesi and Münch, 2020). In particular, the activity of neuroprotective receptors AT1, Sigma-1, and TREM2, as well as induction on Nrf2 pathways in order to secrete neurogenic factors and anti-oxidative enzymes, would be beneficial (Jarrott and Williams, 2016) (Gyengesi and Münch, 2020). Psychedelics are recently reported as agents which display neurogenic, neuroplastic, and anti-inflammatory properties, and they are characterized as physiologically relatively safe agents (Nichols, 2016). Therefore, psychedelics might be an attractive strategy to consider and should be further investigated.

The aim of the present study was to investigate *in vitro* if and how psychedelics can modulate the microglia immune response, which may influence the process of neural tissue cleaning and regeneration. Here we presented results supporting the hypothesis that psychedelic tryptamines (DMT and psilocin) have an immunomodulatory effect on CD11b+ microglia cells, limiting neural cell phagocytosis (psilocin).

Special attention was paid to the expression of hallmark immunoregulatory proteins such as TLR4, Nf-κβ (p65), and CD80 and to recently wider studied TREM2 receptor, which might be a target in the treatment strategy for the neurodegenerative disorders. We observed changes in the protein level of the major markers of the microglial immune response: CD80, TLR4, TREM2, and Nf-κβ on the CD11b+ microglia cells after 100 µM DMT or 100 µM psilocin treatment. Both DMT and psilocin displayed immunomodulatory effects; however, those observed in the psilocin-treated group were stronger on average. In the VEH group, psilocin caused a reduction of CD80 and Nf-κβ (p65) fluorescence intensity by 11-12% more than DMT. In the LPS group, psilocin reduced TLR4 and CD80 fluorescence intensity by 7-10% more than DMT. The difference was observed in the LPS group, where DMT reduced Nf-κβ (p65) fluorescence intensity by 28% stronger than psilocin.

The TLR4 receptor is the component of the innate immunity that reacts to PAMPS such as LPS and DAMPs, (ex. S-100, heat shock proteins, histones, and some other cellular debris) (Roh and Sohn, 2018). TLR4 stimulation activates the NF-κβ, a key factor involved in the inflammatory response acting via the canonical pathway (Liu et al., 2017). It leads to upregulation of co-stimulatory molecules, such as CD80, on the surface of antigen-presenting cells (APC), which is essential to introduce the adaptive immune response into play (Hoebe et al., 2003). Upregulation of co-stimulatory molecule protein level resulting from TLR4 - NF-κβ pathway induction is known as a hallmark response to LPS stimulation in APC cells (Zhang and Ghosh, 2001). The visible decrease in CD80 protein level on microglial cells, both: LPS-stimulated and in the VEH group after psilocin treatment, suggests that it may limit APC ability to present the antigen to T-cells, which might be helpful when considering cellular grafts as a therapeutic strategy. The adaptive immune response is essential for protection against new pathogens. However, its outstanding sensitivity and rapid activation of host immune responses provide a severe limitation when using cell therapy based on brain grafts or transplants (Hoornaert et al., 2017). Therefore, the observed decrease in the CD80 protein expression after psilocin and DMT treatment, together with their reported activity towards induction of neurogenesis and neuroplasticity, is promising (Catlow et al., 2016; Ly et al., 2018).

TREM2 is the transmembrane amine receptor of the immunoglobulin family that regulates inflammatory response and phagocytosis. It is present on tissue-specific macrophages, including microglia and osteoclasts. (Colonna, 2003). The results presented in our study showed that in the VEH group, psilocin increases TREM2 fluorescence intensity on primary microglia by 23% stronger than DMT. LPS treatment increased TREM2 expression on microglia cells by 72% as compared to the VEH group, as LPS is one of the TREM2 ligands. In the LPS group, psilocin increased TREM2 protein expression by 42%, whereas DMT reduced it by 20%. The ability of psilocin to promote upregulation of TREM2 protein level while decreasing pro-inflammatory proteins, even in the situation of LPS-treatment, suggests its potent anti-inflammatory properties. The particular activity of psilocin might be involved in its reported therapeutic effects.

*In vivo*, TREM2 supports microglia-mediated rearrangements of synaptic connectivity in a process called synaptic pruning (Jay et al., 2019) and phagocytosis of apoptotic neurons (Takahashi et al., 2005). Abnormalities in TREM2 regulation might be one of the causes of the progression of diseases such as Alzheimer’s, Parkinson’s, epilepsy, and schizophrenia (Jay et al., 2017). Moreover, the TREM2 expression is involved in phagocytosis, but its exact role in the process is still poorly understood. However, it is already known that TREM2 activation induces DAP12 phosphorylation, leading to cytoskeleton reorganization crucial for phagocytosis. (Takahashi et al., 2005). Interestingly, the abnormalities in TREM2 expression result in dysfunctional phagocytosis of Amyloid β and apoptotic neurons by microglia (Wang et al., 2015) and impairing engulfed particle metabolism (Nugent et al., 2020). It is not completely clear why psilocin but not DMT increased TREM2 protein level in microglia after LPS-stimulation. Both psychedelics share similarities in molecular structure but display significantly varying binding affinities towards receptors [Ray, 2010], resulting in different activation of anti-inflammatory pathways depending on conditions.

In the physiological conditions, miroglia-expressing TREM2 is abundantly localized throughout the brain (Tan et al., 2020). During brain damage, the TREM2 pathway may protect the brain and stimulate the transition of microglia towards anti-inflammatory phenotype (Deczkowska et al., 2020). It was reported in bone-marrow-derived macrophages that TREM2-DAP12 signaling antagonizes TLRs receptors, causing a downregulation in TLRs proteins level and pro-inflammatory cytokine production (Hamerman et al., 2006; Turnbull et al., 2006), whereas silencing of TREM2 attenuated LPS-induced TLR4 signaling and its pro-inflammatory response (Zhong et al., 2015). The downregulation of TLR4 was reported to act protectively for tissues in the model of cerebral ischemia/reperfusion through reduction of inflammatory protein levels (Wang et al., 2014). Therefore, the psychedelic-induced downregulation of TLR4 might be beneficial in brain regeneration.

## Conclusion

The study provides evidence supporting the hypothesis that the DMT and psilocin influence mouse microglia phenotype *in vitro*, and microglia-neural interactions in co-cultures. The psilocin displays more substantial immunomodulatory potential on CD11b+ microglia than DMT, leading to downregulation of pro-inflammatory factors (TLR4, p65, and CD80) and upregulation of TREM2. Psilocin but not DMT also attenuate healthy neurons’ phagocytosis by microglia. Summarizing, DMT and psilocin attenuate microglial pro-inflammatory response and can be considered therapeutic molecules to support neural tissue cleansing and regeneration in multiple conditions with an inflammatory pathogenic component.

## Availability of data and materials

The complete and processed datasets are available along with the manuscript as supplementary material, while the raw data will be available from the corresponding author on request.

## Acknowledgments

We thank friends from the Institute of Immunology and Experimental Therapy Polish Academy of Sciences, especially Dr. Aleksandra Bielawska-Pohl, for the support necessary to finish the project and also: Dr Agnieszka Krawczenko, Dr Maria Paprocka, M.Sc. Agnieszka Szyposzyńska, M.Sc. Elżbieta Wojdat and M.Sc. Olga Doszyń for help and advices in lab. We also thank Institute’s handymen for adjusting the store room to meet the requirements for psychedelic drug storage. We also thank Dr. Andrzej Żak from the Wroclaw University of Technology for his friendly energy and words of support.

## Abbreviations

APC: Antigen Presenting Cell
AD: Alzheimer’s Disease
BBB: Blood-Brain Barrier
CSAIDs: cytokine-suppressive anti-inflammatory drugs
CTCF: The Corrected Total Cell Fluorescence
DAMPS: Danger Associated Molecular Patterns
FCM: flow cytometry
IF: immunofluorescence
LPS: lipopolysaccharide
NPC: Neural Progenitor Cells
PAMPs: Pathogen Associated Molecular Patterns
PTSD: Post Traumatic Stress Disorder
TBI: Traumatic Brain Injury
TLR4: Toll-like Receptor 4
TRD: Treatment-Resistant Depression.

## Author contributions statement

UK conceived, designed, supervised all experiments, analyzed the data and wrote the manuscript. UK was responsible for the research concept and obtaining funding. AK provided scientific guidance, KW obtained NPC cells, and helped with the manuscript writing and editing process. MF analyzed the data, wrote and edited the manuscript, provided scientific guidance, and part of the funding.

## Ethics Statement

DMT and psilocin were obtained legally after acquiring permission from the Pharmaceutical Inspectorate in Wroclaw city, Poland (WIF-WR I.857.6.2.2020).

## Figure Legend

**Fig S1: Characteristics of the brain isolated E10 embryonic bodies differentiation:** The cells were characterized with an ability to differentiate in 7 days in DMEM/F12 supplemented with 1% B27, 1% Pen/Strep, 1% L-glutamine, 50 ng/mL BDNF, and 30 ng/mL GDNF into cells displaying neural morphology (A). Positive immunofluorescent staining with Synapsin and β-III-tubulin confirmed successful neural differentiation (B). In the differentiated neuronal culture, there was no GFAP-positive fraction; however, the population of single microglial cells could be identified with IBA-1+ staining(arrows). Control staining revealed no unspecific reactions (E), scale bar = 50µm.

**Fig. S2: Methodology of Corrected Total Cell Fluorescence (CTCF) measurement**.

**Fig. S3: The DMT and psilocin 12h stimulation decrease fluorescence intensity of TLR4 on CD11b+ microglia cells after LPS treatment (double staining)**. This figure is related to Fig. 4. To identify microglia, the cells were double stained with CD11b marker (Alexa fluor 488nm).

**Fig. S4: DMT and psilocin downregulate Nf-**κβ **p65 protein level in both control and LPS conditions (double staining)**. This figure is related to Fig. 5. To identify microglia, the cells were double stained with CD11b marker (Alexa fluor 488nm).

**Fig. S5: Psilocin but not DMT upregulates TREM2 protein level in VEH and LPS-stimulated group (double staining)**. This figure is related to Fig. 7. To identify microglia, the cells were double stained with CD11b marker (Alexa fluor 488nm).

