## Supplementary figures and images for "The DMT and Psilocin Treatment Changes CD11b+ Activated Microglia Immunological Phenotype"

### Supplemental Figure 1

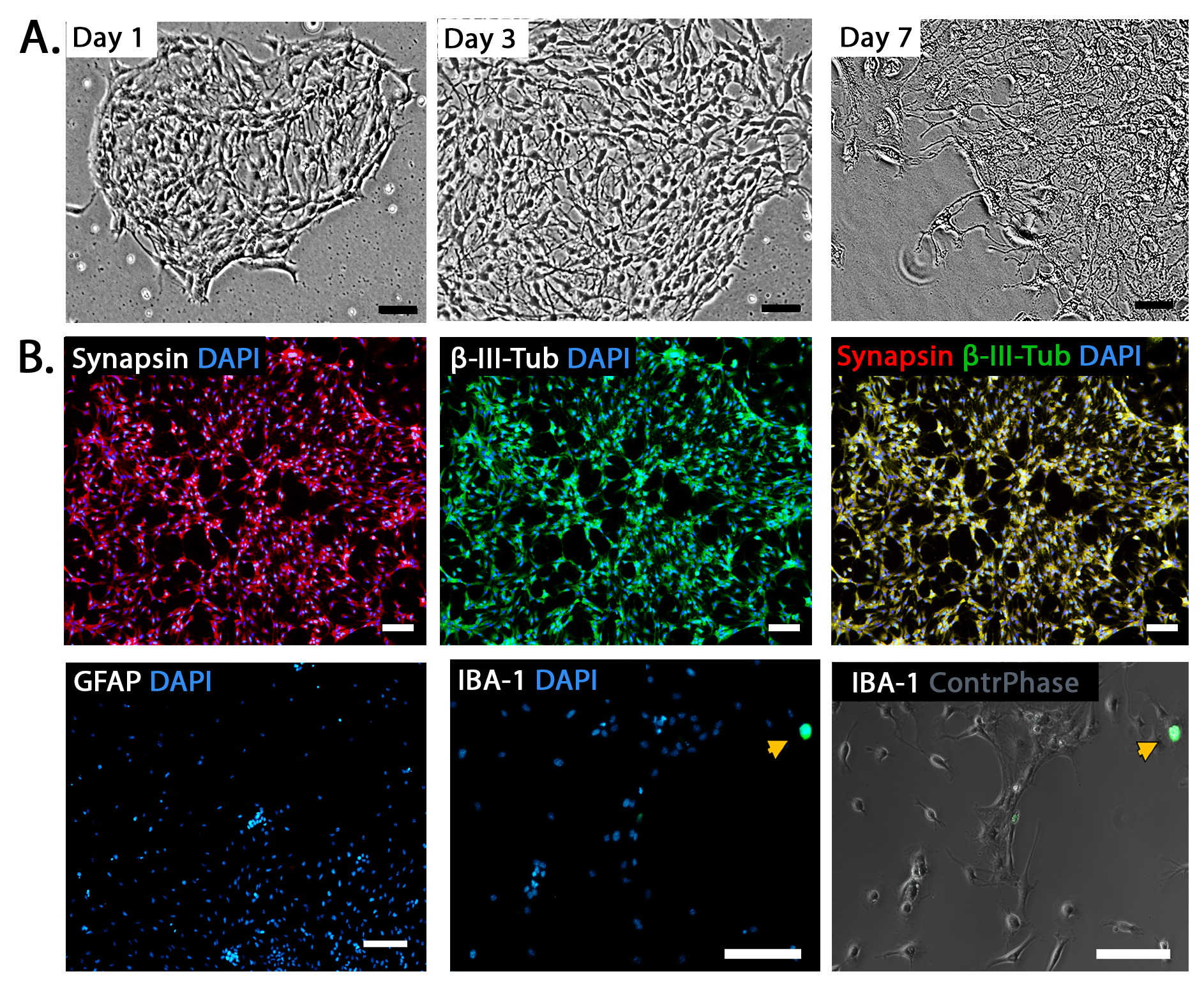

### Supplemental Figure 2

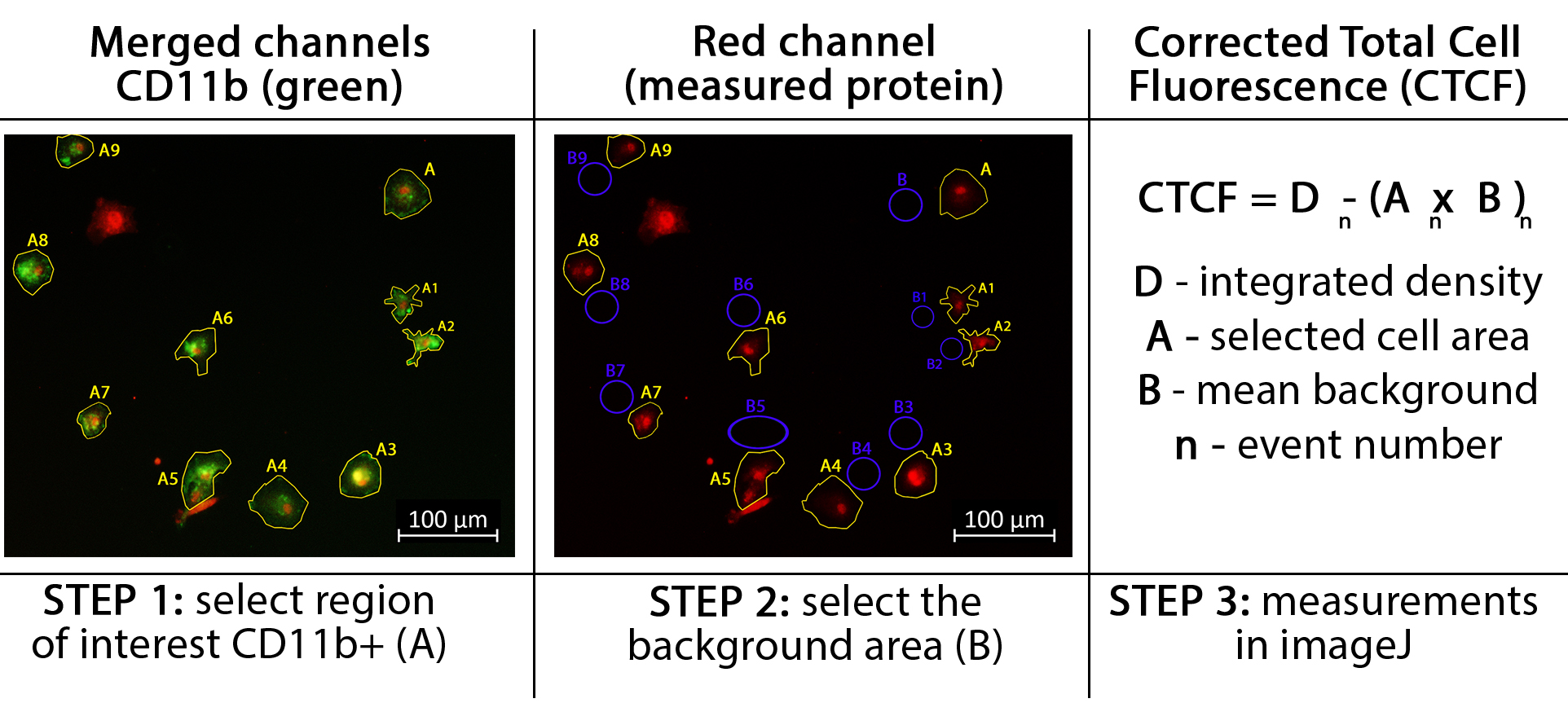

### Supplemental Figure 3

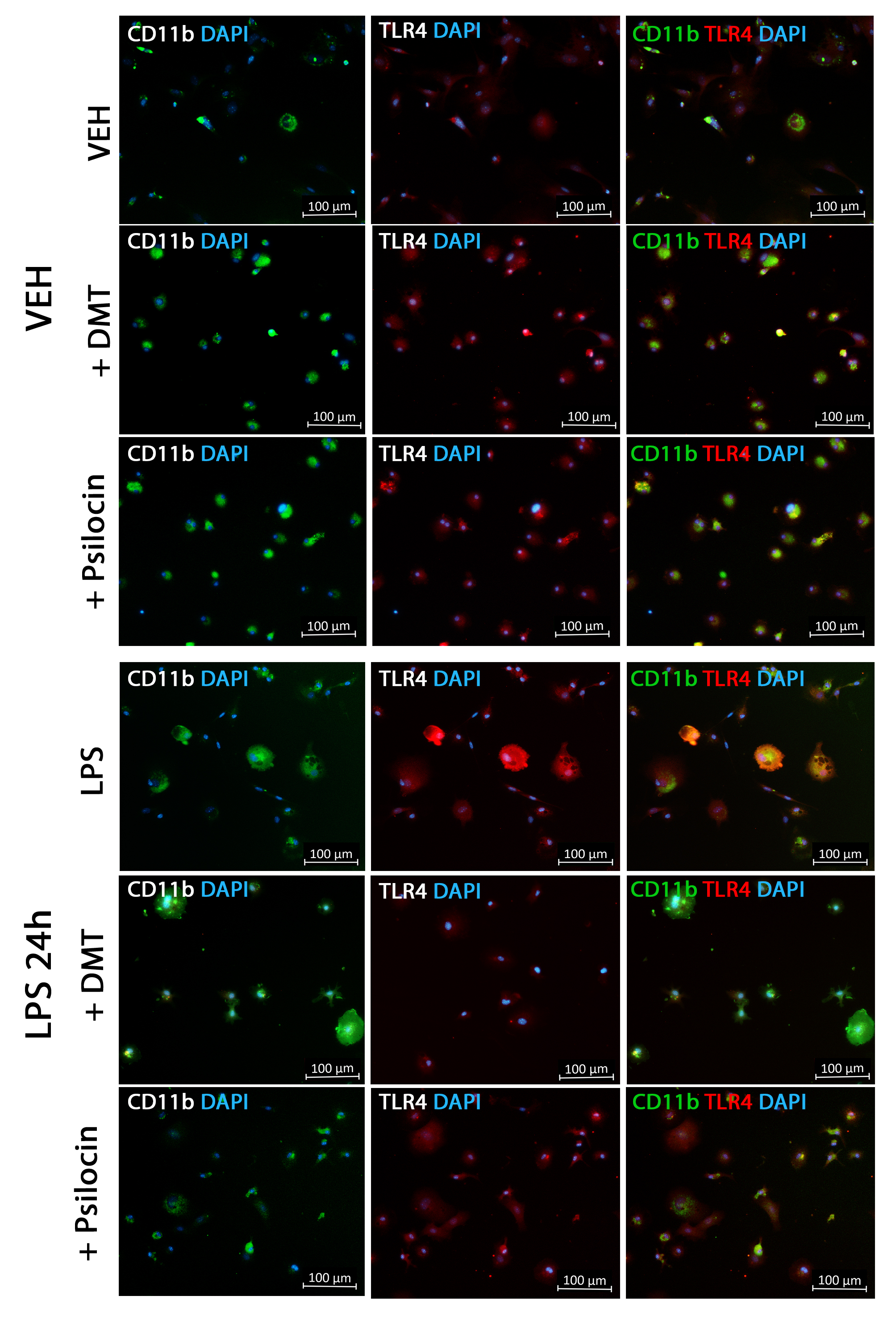

### Supplemental Figure 4

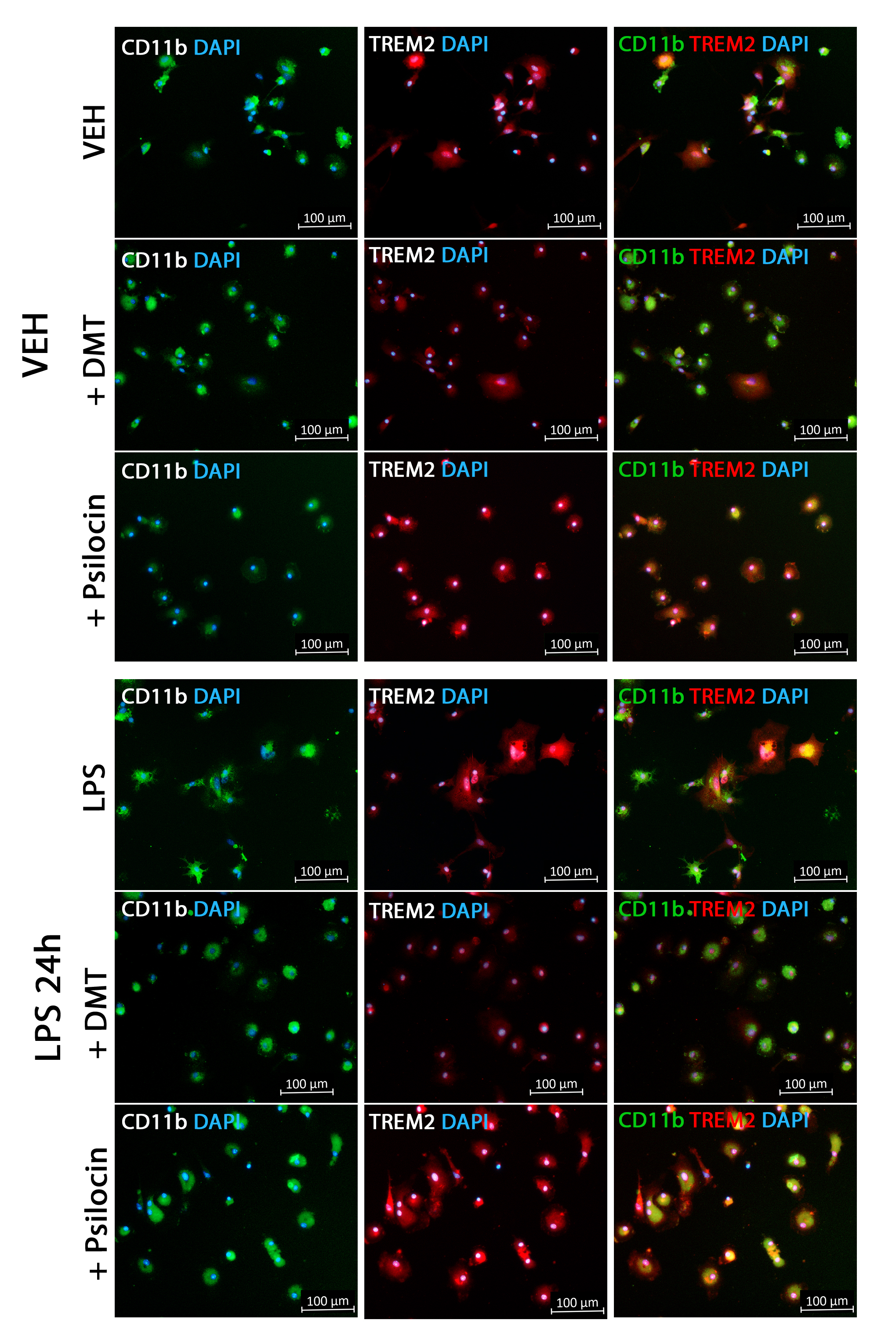

### Supplemental Figure 5

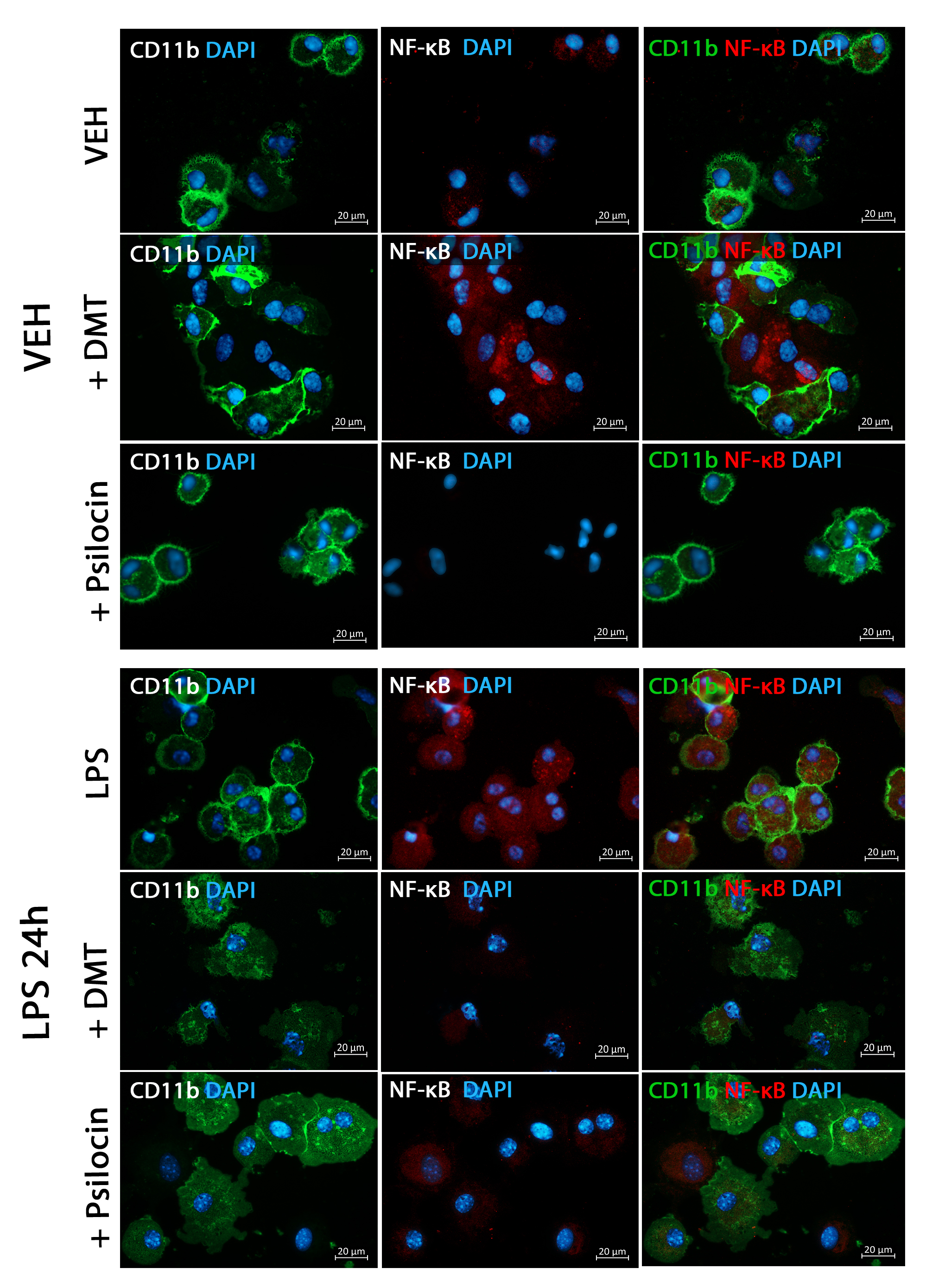
